# How many segments are enough to biomechanically model the feet? A comparison of inverse kinematics and dynamics in multisegmented foot models

**DOI:** 10.1101/2024.05.13.593935

**Authors:** Julia Nicolescu, Logan Gaudette, Olivia Vogel, Irene S. Davis, Adam S. Tenforde, Karen L. Troy

## Abstract

Multisegmented foot models (MSFMs) capture kinematic and kinetic data of specific regions of the foot instead of representing the foot as a single, rigid segment. Models differ by the number of segments and segment definitions, so there is no consensus for best practice. It is unknown whether MSFMs yield the same joint kinematic and kinetic data and what level of detail is necessary to accurately measure such values. We compared the angle, moment, and power measurements at the tibiotalar, midtarsal, and metatarsophalangeal joints of four MSFMs using motion capture data of young adult runners during stance phase of barefoot walking and jogging. Of these models, three were validated: Oxford Foot Model, Milwaukee Foot Model, and Ghent Foot Model. One model was developed based upon literature review of existing models: the “Vogel” model. We performed statistical parametric mapping comparing joint measurements from each model to the corresponding results from the Oxford Model, the most heavily studied MSFM. We found that the Oxford Foot Model, Milwaukee Foot Model, Vogel Foot Model, and Ghent Foot Model do not provide the same results. The changes in model segment definitions impact the degrees of freedom in ways that alter the measured kinematic function of the foot, which in turn impacts the kinetic results. We also found that dynamic function of the midfoot/arch may be better captured by MSFMs with a separate midfoot segment. The results of this study capture the variability in performance of MSFMs and indicate a need to standardize the design of MSFMs.

## Introduction

The human foot is complex, yet it is often represented as a single segment in biomechanical modeling. Such models are unable to capture the kinematic and kinetic data of specific regions of the foot. To understand injuries in the feet, such as bone stress injuries, plantar fasciitis, Achilles tendonitis, and arthritis, a multisegmented foot model is necessary. Multisegmented foot models (MSFMs) have become a core part of extracting biomechanical information in research and in the clinic (Leardini et al., 2019). These models allow researchers to observe kinematic and kinetic behavior at specific joints within the feet. Our lab is interested in the tibiotalar, midtarsal, and the metatarsophalangeal joints as they relate to metatarsal bone stress injuries in long-distance runners. It has been shown that midfoot motion during gait is a relevant factor that should be accounted for in kinematic foot research (Bassett et al., 2023). By utilizing a MSFM instead of treating the foot as a rigid segment, we were able to explore midfoot motion using the kinematics and kinetics of joints proximal and distal to the metatarsals in addition to the ankle.

Across the various MSFM that have been developed, each differs by the number of segments included and how they are defined, so there is no consensus for best practice. A recent review by Leardini et al. documents the various MSFMs currently available, their number of segments, and whether they have been validated (Leardini et al., 2019). Two widely used and validated four-segment models are the Oxford Foot Model and the Milwaukee Foot Model (Carson et al., 2001; Sampath et al., 1998). Each of these models contain shank, rearfoot, forefoot, and hallux segments. A key difference between the models is that the Oxford Foot Model excludes the midfoot/tarsals from the forefoot, leaving a gap between the rearfoot and forefoot. The Milwaukee Foot Model includes the midfoot/tarsals in the forefoot section, eliminating the break featured in the Oxford Foot Model. (Carson et al., 2001; Sampath et al., 1998). Models increase in complexity as the number of segments and degrees of freedom increase. Oosterwaal et al. even implemented a 26-segment foot model (Oosterwaal et al., 2011). Yet it is important to note that as more segments are introduced, more assumptions must be made. Additionally, the more segments there are, the smaller the segments become. These small segments are difficult to track using optical motion capture techniques which is a common approach for MSFMs. Other models achieve higher levels of detail by adding a medial/lateral split within the forefoot, like the Ghent Foot Model (De Mits et al., 2012). Thus, the first step to implementing a MSFM in research or in clinic is attempting to identify one out of the many available that will be best suited for the work at hand.

It is unknown whether the various MSFMs available yield the same joint kinematic and kinetic data and what level of detail is necessary to accurately measure such values. Kinematic studies comparing available MSFMs have been completed and variation between models has been reported (Nicholson et al., 2018; Powell et al., 2013). Nicholson et al., reported that between the five MSFMs analyzed, patterns of movement were similar, but there were frequent offsets between models that need to be considered when using data published from these models. Powell et al., suggested that both the MSFMs used in their study detected change in frontal plane motion between low- and high-arched athletes, but that one of the models, the Leardini model, may be more appropriate because it can track midfoot motion during dynamic motion while the Oxford Foot Model cannot. There is a lack of information regarding the kinetic results of MSFMs and very few comparing kinetic information. **This is important because kinematic and kinetic information about the joints within the foot enables us to understand foot posture and function in relation to injury risk**. To our knowledge, existing works do not address the assumptions and definitions of the MSFMs that may contribute to variation in model results. It is essential to explore the features of MSFMs that may influence the joint kinematic and kinetic measurements to identify a best practice for MSFMs.

To address these uncertainties, we compared angle, moment, and power measurements at the tibiotalar, midtarsal, and metatarsophalangeal joints of four MSFMs using motion capture data of young adult runners during stance phase of barefoot walking and jogging. Of these models, three were validated MSFMs: the Oxford Foot Model, the Milwaukee Foot Model, and the Ghent Foot Model. One model was developed in our lab based upon literature review of existing models, referred to as the “Vogel” model. By using the same input data for each model, we were able to examine how the assumptions and segments defined in each MSFM may influence the joint kinematic and kinetic results obtained. Tibiotalar, midtarsal, and metatarsophalangeal joint angle, moment, and power were quantified in the sagittal plane and the intermodal differences were compared. We hypothesized that increasing the number of model segments would decrease the power and moment at individual joints because it will be spread across the various segments. Simultaneously, we expected that increasing the number of model segments would lead to more variation in the results due to the increasing degrees of freedom and increasing need for assumptions within the model. We expect the results for the tibiotalar joint to be similar across all the models based on the similarities in the shank and rearfoot segments in each model.

## Methods

### Participants/Study Design

The criteria for participation in this study were adults aged 18-30 who ran an average of at least 24 km per week during the last 3 months. We recruited 20 young adults to participate in this institutionally approved study with an average weekly running volume of 41.4 + 7.7 km. All participants gave written informed consent to participate in this institutionally approved study. Complete participant demographics are summarized by Table 1.

**Table 1.** Participant demographics: sex, age, mass, height, and weekly running volume.

|  | Male | Female | Whole Group |
| --- | --- | --- | --- |
| **Participants** | 10 | 10 | 20 |
| **Age (years)** | $21.8 \pm 1.3$ | $22.0 \pm 1.3$ | $21.9 \pm 1.3$ |
| **Mass (kg)** | $66.9 \pm 6.0$ | $57.4 \pm 5.0$ | $62.1 \pm 7.3$ |
| **Height (m)** | $1.8 \pm 0.1$ | $1.7 \pm 0.0$ | $1.7 \pm 0.1$ |
| **Weekly Running Volume (km)** | $44.4 \pm 6.4$ | $37.5 \pm 7.3$ | $41.4 \pm 7.7$ |

### Motion capture

Motion and force data were collected with a 10 camera Vicon Vantage system (Vicon, UK) operating at 100 Hz, and two 6-degree of freedom force platforms embedded in the laboratory floor (AMTI, Watertown) operating at 1,000 Hz. The marker set used during this study was based upon the suggested markers for the MSFM designed at KU Leuven (Malaquias et al., 2017). This consisted of 16 markers on each foot and 8 on each shank with a total of 52 markers. Markers had radii of 7 or 4.5 mm and were placed according to bony landmarks when possible, and otherwise placed anatomically. This study only considered the foot and shank, and the marker placement for these segments is shown in Figure 1. A review of marker sets for the various MSFMs used in this study showed that our set was more comprehensive (Carson et al., 2001; De Mits et al., 2012; Sampath et al., 1998). It is important to note that this marker set was used to construct all the models included in this work, and that the original published and validated models do not include all the same markers. A comparison of the number and location of markers used to define specific segments in each model is shown in Appendix A.

**Figure 1.**
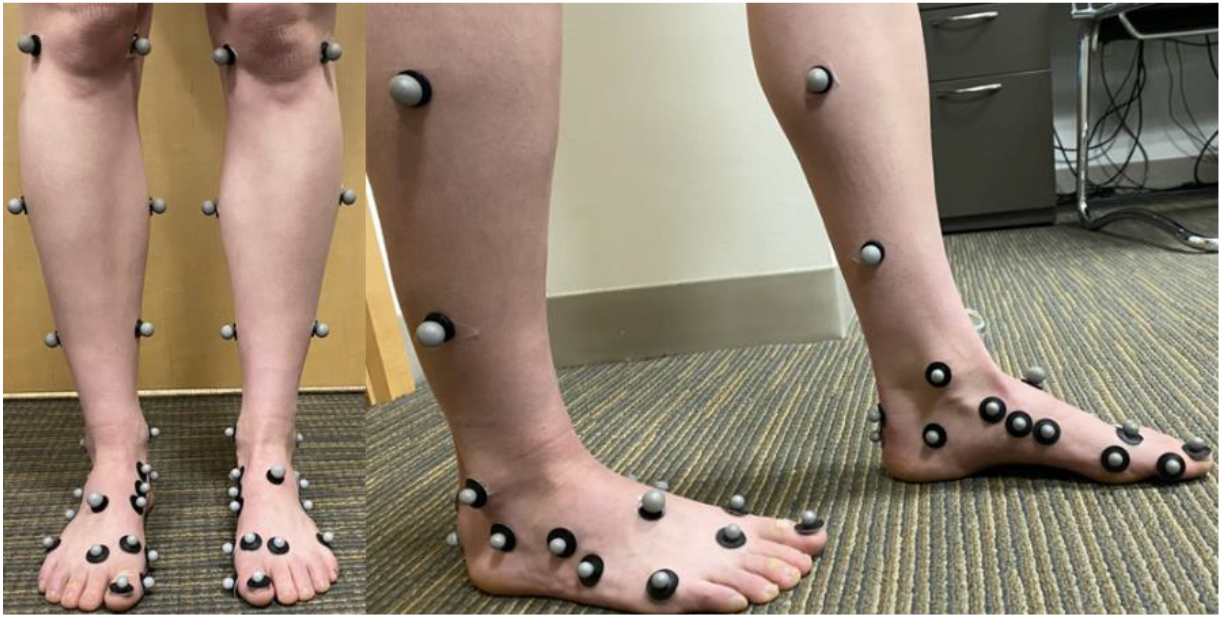
Foot and shank marker placement.

Participants performed barefoot walking and jogging on a 10-meter runway with two force plates embedded at the center. During trials they were instructed to look ahead rather than at the floor to avoid intentional targeting of the force plates. The walking trials were performed at a self-selected speed. The slow jog was instructed to represent a warmup jog speed. Trials were repeated until the participant’s left and right feet each landed fully on the force plates.

Trials were excluded for analysis if the ground force reaction vector was assigned erroneously outside of the area of the foot contacting the force plate surface. For this study, only data for the right foot was included due to a technical issue with the left force plate. Manual labeling and gap filling was performed using Vicon Nexus software (Version 2.15.0) for all trials. Analog data and trajectories were filtered using fourth order Butterworth filters with a cutoff frequency of 300 Hz and 6 Hz respectively. Automatic gap filling was done using spline fill or pattern fill when manual gap filling was not an option. Complete data were available for 18 of the 20 participants.

### Model Selection

We performed a literature review to identify available MSFMs and the features of each one. The Oxford Foot Model was selected as the primary reference point for comparison due to its widespread use and validation. The Milwaukee Foot Model was selected because it is similar to the Oxford Foot Model but offers slight variation in the midfoot section. It includes the midfoot region as a part of the forefoot segment while Oxford excludes the midfoot from the forefoot, leaving a break in between the rearfoot and forefoot. We designed the Vogel Model after extensive literature review of the existing MSFMs because we saw that no current model included a segment containing all the phalanges, and few models included a separate midfoot section. It includes a separate midfoot segment and a phalanges segment (as opposed to hallux segment). Lastly, the Ghent model was selected because it introduced a medial/lateral split and a separate midfoot section. Figure 2 shows the anatomical segments of each model. Because our interest is in understanding bone stress injuries of the metatarsals, we only selected MSFMs that contained a segment starting or ending at the metatarsals. Each model was constructed using Visual 3D software (C-Motion, Inc., Maryland). The number of segments and segment definition of each MSFM is shown in Table 2 and illustrated in Figure 2.

**Table 2.** The MSFMs included in the study, segment number, segment definition, and how they were designed in Visual3D software (C-Motion, Inc., Maryland).

| Model Name | Number of Segments | Segments | Construction in Visual 3D | Reference |
| --- | --- | --- | --- | --- |
| **Oxford Foot Model** | 4 | Shank, rearfoot, forefoot (excluding distal tarsals), hallux | Adapted from C-Motion tutorial | (Carson et al., 2001; C-Motion, 2018) |
| **Milwaukee Foot Model** | 4 | Shank, rearfoot, forefoot (including distal tarsals), hallux | Adapted from Oxford | (Sampath et al., 1998) |
| **Vogel Foot Model** | 5 | Shank, rearfoot, midfoot, forefoot, phalanges | Original construction | Appendix B |
| **Ghent Foot Model** | 6 | Shank, rearfoot, midfoot, medial forefoot, lateral forefoot, hallux | Description of construction from original authors | (De Mits et al., 2012) |

**Figure 2.**
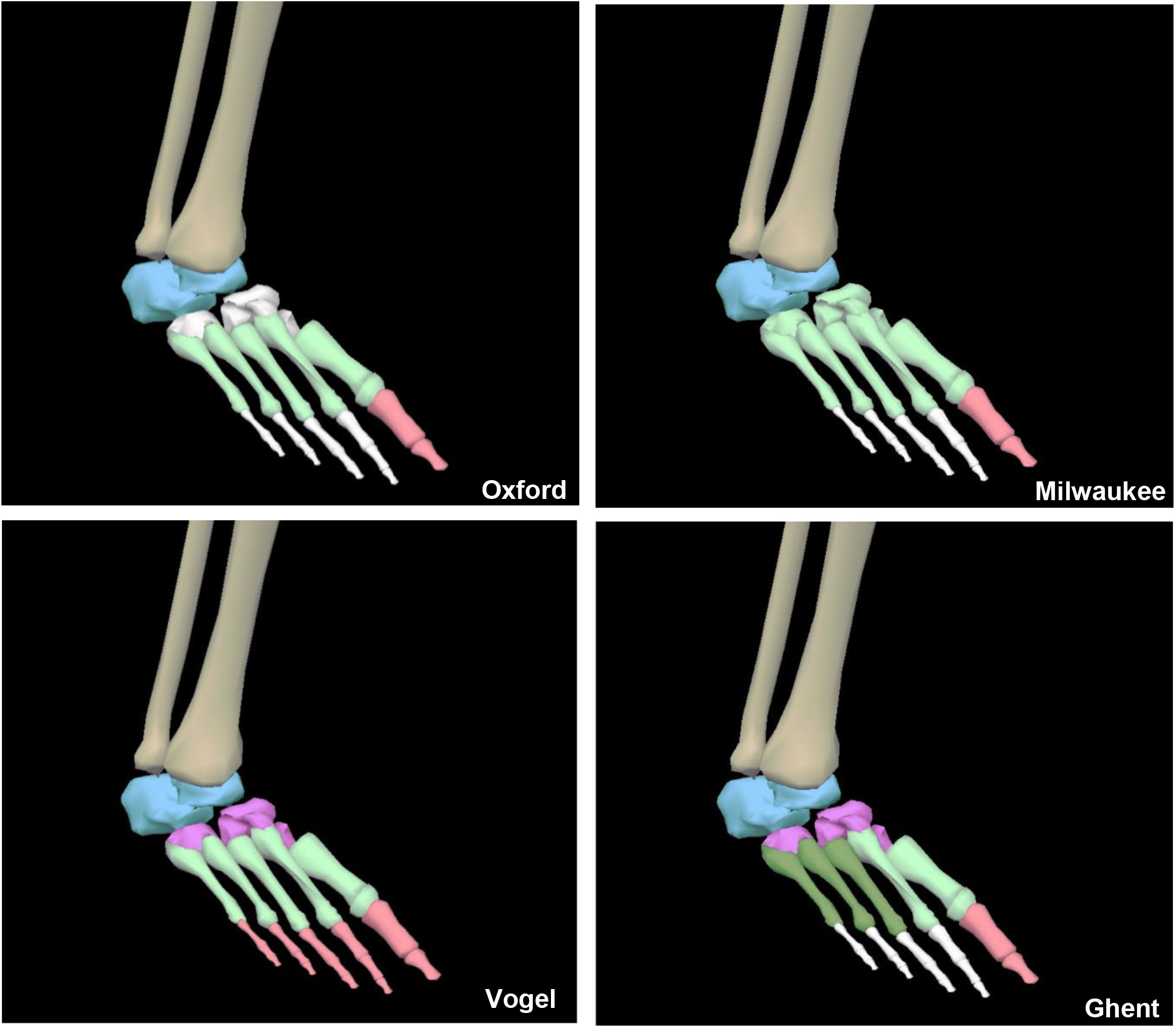
Oxford, Milwaukee, Vogel, and Ghent foot models with model segments. White bones indicate that they were not included in the model segments.

A pipeline was written to calculate the joint angle, moment, and power of the tibiotalar, midtarsal, and metatarsophalangeal joints during each trial for each model. For angle measurements, the angle was in reference to the proximal segment. The joint moment and power were resolved around the laboratory coordinate system where the positive y-axis is coronal, x-axis is sagittal, and the z-axis is axial. Only the stance phase was analyzed here, expressed as a percentage (1-100%) to facilitate comparisons between participants and models.

### Statistical Analysis

In total, we calculated three parameters (angle, moment, power) at three joints (tibiotalar, midtarsal, and metatarsophalangeal) for four models (Oxford, Milwaukee, Ghent, and Vogel) across two speed conditions (walking, jogging). For each condition, we calculated means and standard deviations of the entire stance phase (1-100%). To test our hypothesis about the effects of adding model segments, we performed statistical parametric mapping (SPM), using a paired t-test statistic (Pataky, 2010). Each model was compared to the Oxford Foot Model, which we considered the primary reference point.

## Results

### Tibiotalar Joint

Our hypothesis that the tibiotalar joint would be similar across each of the models was not entirely supported. Although model kinematics were similar during walking, tibiotalar moments were significantly greater at the end of the stance phase for each model compared to the Oxford model (Figure 3). The Oxford to Vogel models had similar tibiotalar power, while the Milwaukee and Ghent models had significantly greater power around 83-92% of stance.

**Figure 3.**
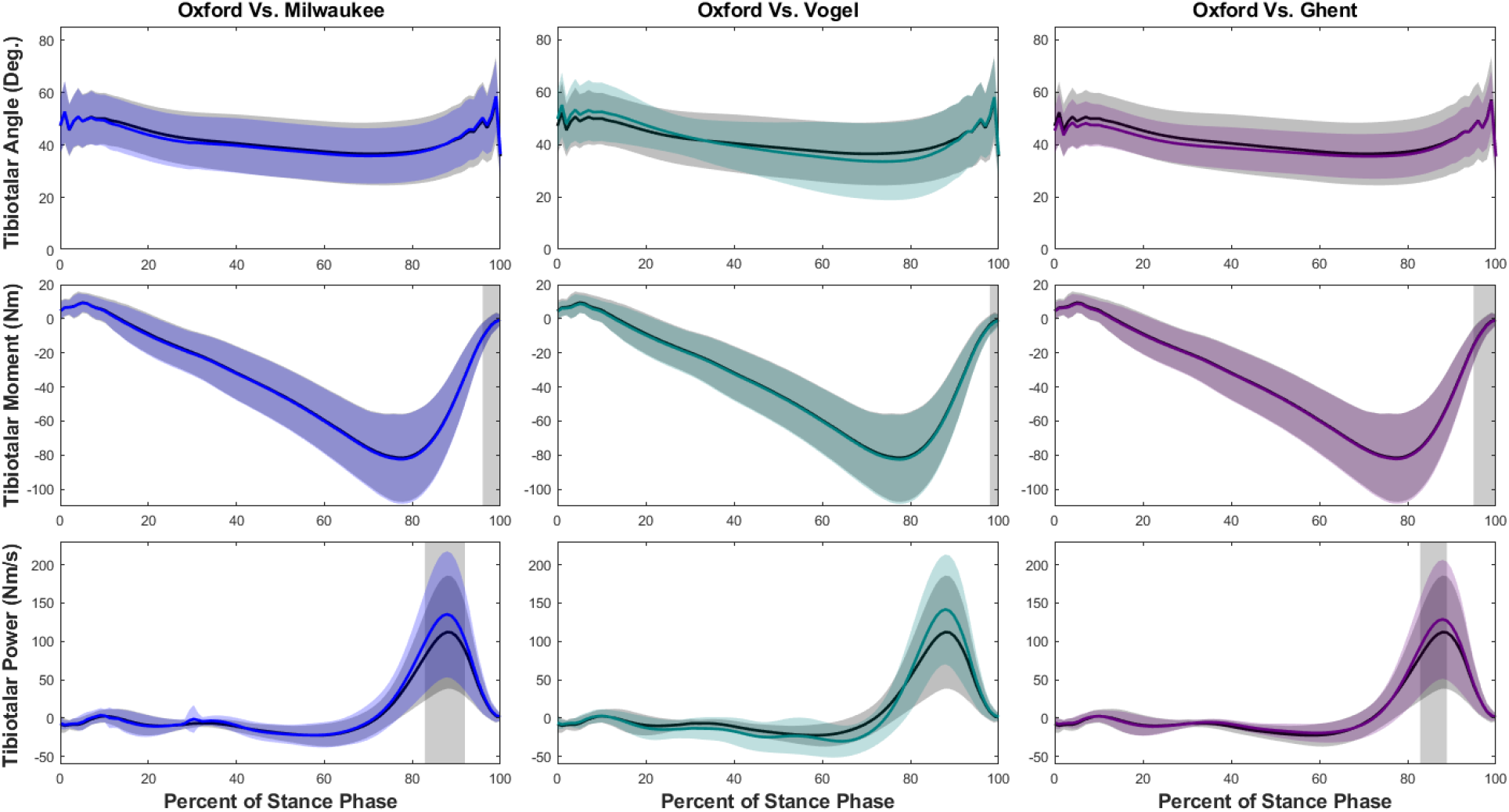
Tibiotalar joint ankle, moment, and power means and standard deviations during walking with SPM results. Oxford is shown in black, Milwaukee is shown in blue, Vogel is shown in green, and Ghent is shown in purple. The grey rectangles show the regions of statistically significant difference between model results.

Tibiotalar joint measurements were less similar between models during jogging (Figure 4). Although the angles were similar between Oxford and Ghent models, the Milwaukee and Vogel models showed a decrease during early to mid-stance. Tibiotalar joint moment was significantly greater in magnitude in each of the models at both mid- and terminal stance when compared to the Oxford model. Tibiotalar joint power was also significantly greater in each model, especially during the second half of stance.

**Figure 4.**
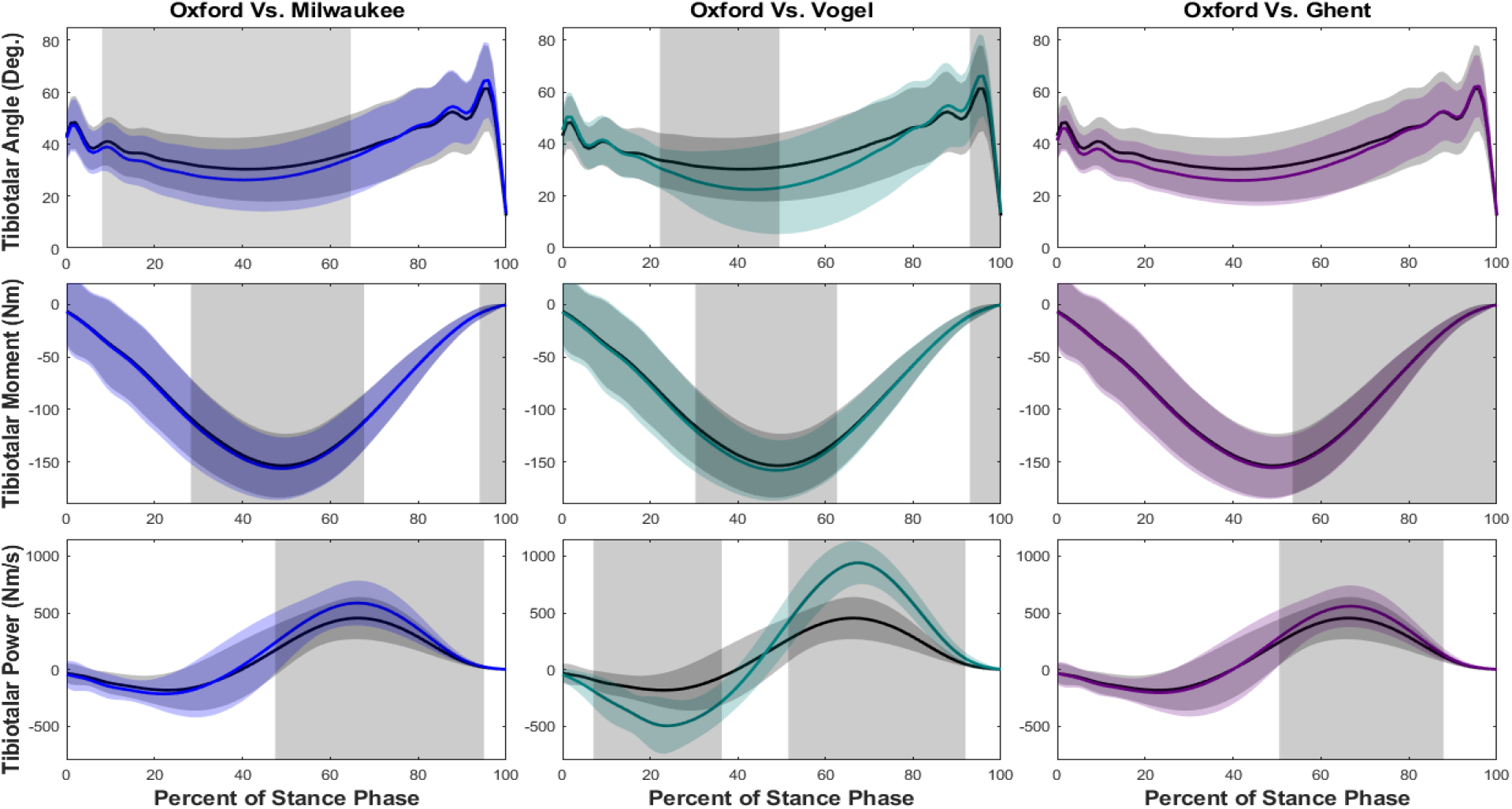
Tibiotalar joint ankle, moment, and power means and standard deviations during jogging with SPM results. Oxford is shown in black, Milwaukee is shown in blue, Vogel is shown in green, and Ghent is shown in purple. The grey rectangles show the regions of statistically significant difference between model results.

### Midtarsal Joint

At the midtarsal (MT) joint during walking (Figure 5), we observed that there were significant differences in the kinematics for the entirety of stance phase in Milwaukee and Ghent compared to Oxford, but no difference between Vogel and Oxford. During jogging (Figure 6) we observed that there were significant differences in the kinematics for almost the entirety of stance phase in all models compared to Oxford. All models showed a decrease in joint angle compared to the Oxford model, although the shapes of the time vs. angle curves were similar. The joint moment was greater in both the Milwaukee and Vogel models for the majority of stance phase in both walking and jogging. However, the Ghent model showed decreased joint moments for most of walking and jogging stance. All three models showed reduced MT joint power at various points, especially in the second half of stance. However, there was no significant difference between Vogel and Oxford MT joint power during walking.

**Figure 5.**
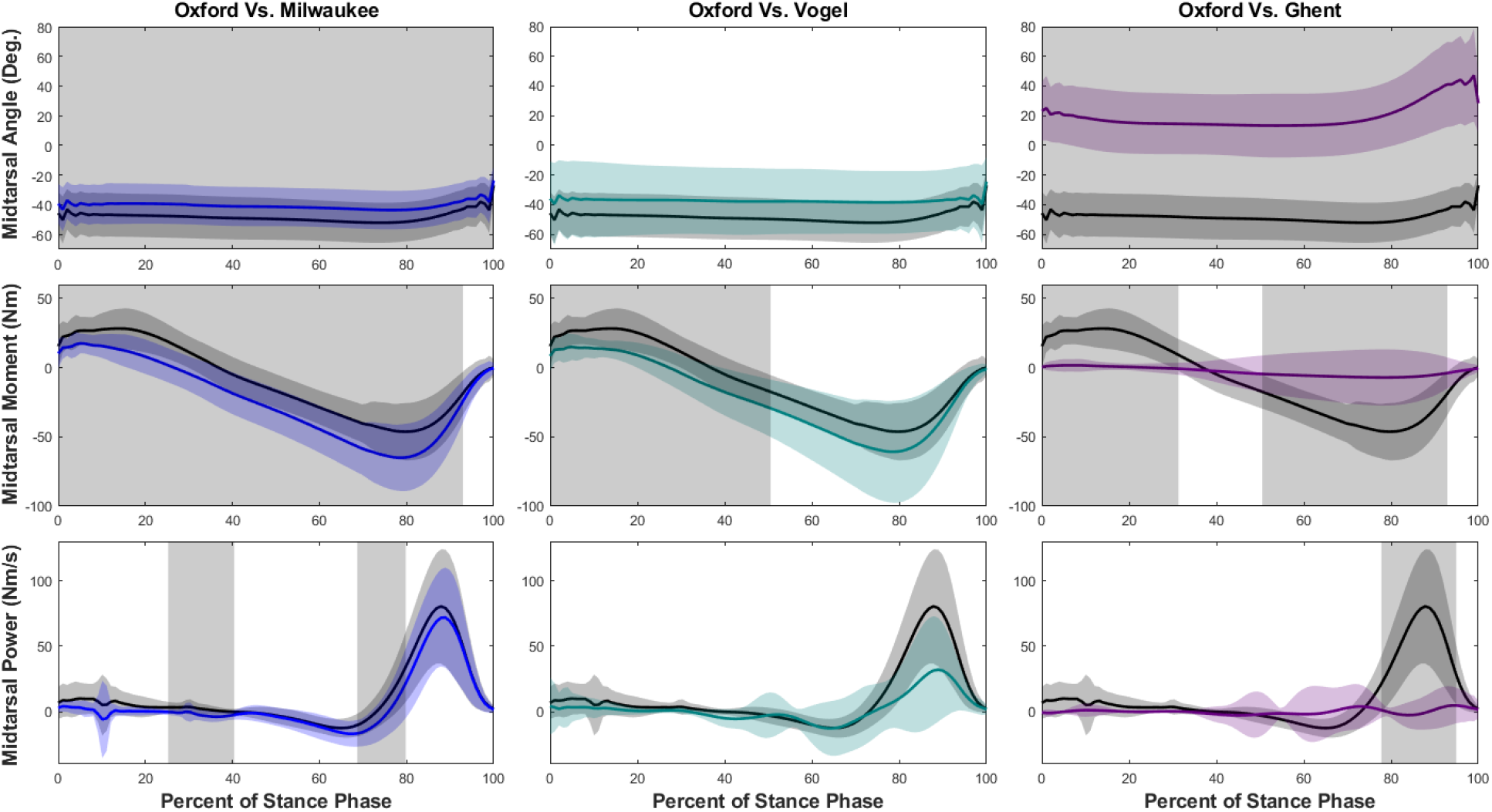
Midtarsal joint ankle, moment, and power means and standard deviations during walking with SPM results. Oxford is shown in black, Milwaukee is shown in blue, Vogel is shown in green, and Ghent is shown in purple. The grey rectangles show the regions of statistically significant difference between model results.

**Figure 6.**
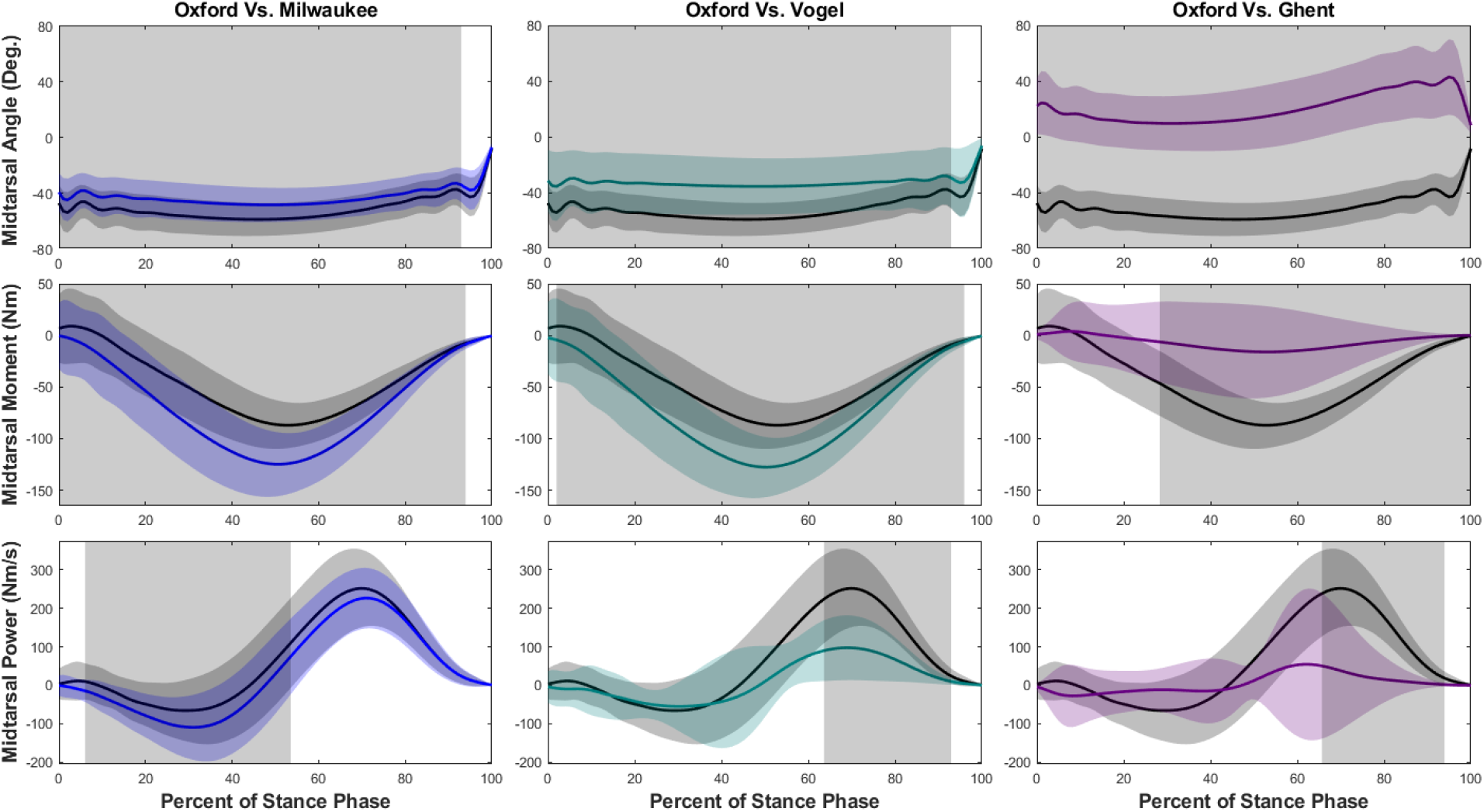
Midtarsal joint ankle, moment, and power means and standard deviations during jogging with SPM results. Oxford is shown in black, Milwaukee is shown in blue, Vogel is shown in green, and Ghent is shown in purple. The grey rectangles show the regions of statistically significant difference between model results.

### Metatarsophalangeal Joint

At the metatarsophalangeal (MTP) joint during walking (Figure 7) and jogging (Figure 8) the kinematic results differed depending on the gait condition. The MTP angle vs. time curves were similar for the Vogel and Milwaukee but were both offset from the Oxford model. In contrast, the Ghent model showed an opposite motion. The MTP moment and power were nearly zero for all models except the Ghent model, which displayed a large moment and significant power in late stance phase.

**Figure 7.**
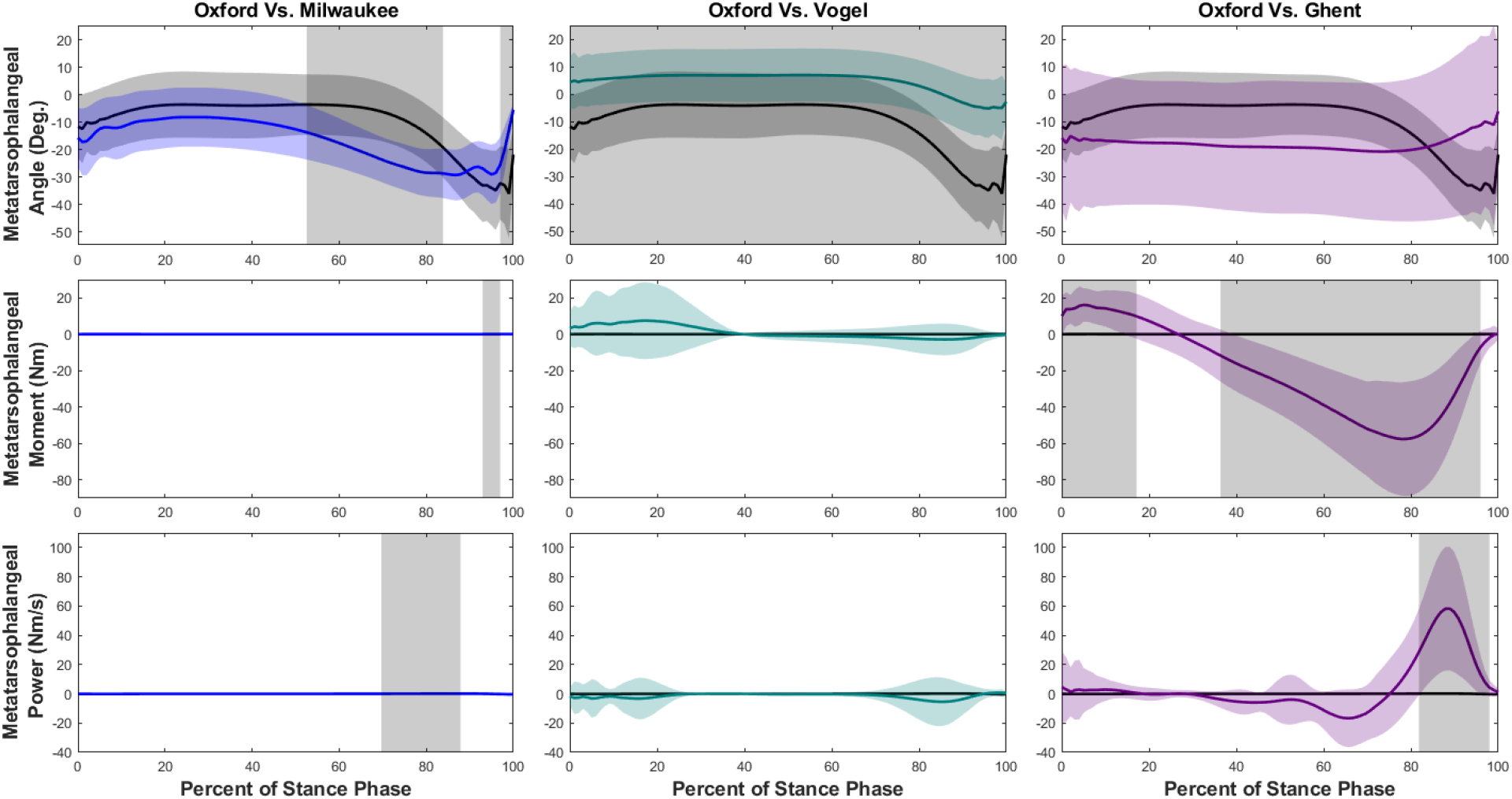
Metatarsophalangeal joint ankle, moment, and power means and standard deviations during walking with SPM results. Oxford is shown in black, Milwaukee is shown in blue, Vogel is shown in green, and Ghent is shown in purple. The grey rectangles show the regions of statistically significant difference between model results.

**Figure 8.**
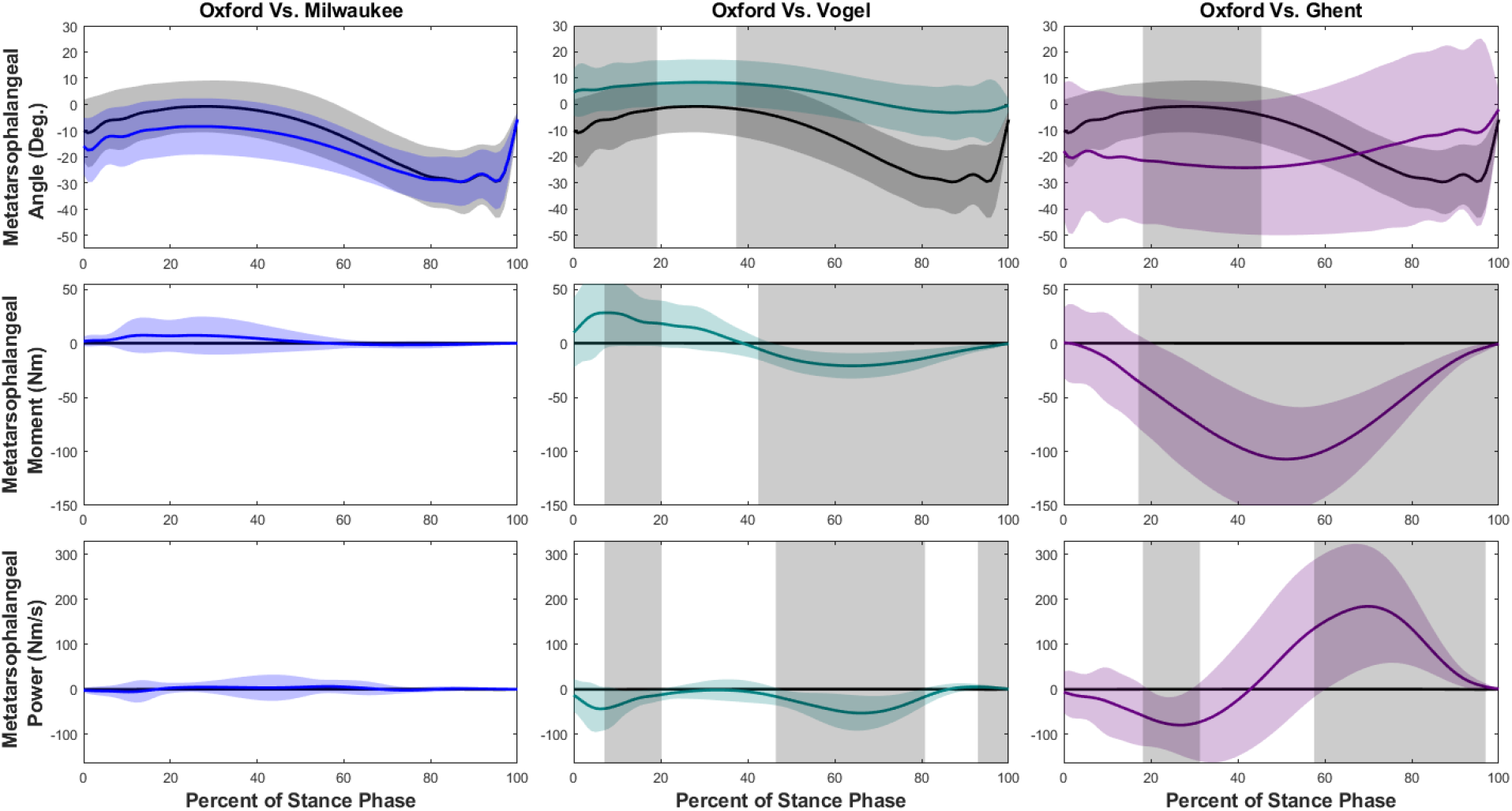
Metatarsophalangeal joint ankle, moment, and power means and standard deviations during jogging with SPM results. Oxford is shown in black, Milwaukee is shown in blue, Vogel is shown in green, and Ghent is shown in purple. The grey rectangles show the regions of statistically significant difference between model results.

## Discussion

Here, we compared the kinematics and kinetics of four MSFMs. Our results demonstrate that even small changes in segment definitions of multisegmented foot models may impact the kinematic and kinetic information of foot joints represented in these models. We originally expected to observe similar results at the tibiotalar joint across the models, based on the similarities in how the shank and the rearfoot are defined. Despite the shared features of the models, the tibiotalar joint moments during walking were different for each of the models during toe-off. During this period, the ankles of most participants were plantarflexed. So, the models were able to calculate comparable results for joint moment for most of stance until a period of high plantarflexion. Moreso, the models generally showed a greater magnitude of tibiotalar moment and power during both walking and jogging when compared to Oxford implying that the exclusion of the midfoot may specifically impact kinetic calculations.

The midtarsal joint was of special interest to us because of its relevance in arch/midfoot motion as it relates to metatarsal bone stress injuries. An original goal of this project was to use a MSFM to quantify midtarsal and tarsometatarsal (TMT) joint biomechanics, which prompted the question of what model should be used. The present results show that MT kinematics and kinetics differ significantly depending on the model. We observed that at the midtarsal joint, the Vogel model performed most similarly to the Oxford model. Even so, there was a large decrease in the midtarsal angle using the Vogel and Ghent models which could be explained by the small size of the midfoot section. A smaller midfoot would likely generate less angular movement between the rearfoot and the midfoot than angular movement between the rearfoot and forefoot segments in the Oxford model. This also explains the smaller magnitude of power seen in these models as less angular acceleration will be observed. This may be an indication of the reality that smaller segments are difficult to track in motion capture. These findings also indicate that models neglecting the midfoot may attribute more power at these joints than what is really occurring.

This means the dynamic function of the midfoot and arch seen in walking and jogging, like arch stiffening or lack thereof occurring at the midfoot, may not be effectively captured by the Oxford model. This notion supports previous findings by Powell et al. who suggested that the Oxford model is not able to track midfoot motion dynamically in comparison to a model with a midfoot segment (Powell et al., 2013). Additionally, if the Vogel model is used for future work, the separate midfoot section allows kinetic and kinematic measurements at the tarsometatarsal joint in addition to the midtarsal joint. We believe that this feature will allow us to better represent the finer movements of the midfoot segment and its role in dynamic arch function. Likewise, other MSFMs with a separate midfoot segment like the Leardini or Rizzoli models provide these benefits as well (Leardini et al., 1999, 2007). Given its kinematic and kinetic validity along with the midfoot segment, we believe the Vogel model will be useful in future work on metatarsal bone stress injuries.

Our hypothesis that increasing the number of model segments would decrease the power and moment at individual joints was not consistently supported. The results of this study are an indication that the variation in the segment definitions has unique impacts on the kinematic data that will also impact the kinetic data. These differences are likely the result of how the kinematics of the segments change directly with alterations in segment definitions. For example, in the Oxford, Vogel, and Ghent models the forefoot segment consists of the metatarsals while the Milwaukee forefoot segment includes the metatarsals and the distal tarsals. One impact of this small change is simply a larger forefoot segment for the Milwaukee model. This leads to differences in how modeling software such as Visual 3D automatically assigns forces to segments, since the assignment relies on the location of the center of pressure in relation to the segments. Greater precision could be achieved with pressure-sensing insoles or a pressure mat to proportionally assign the ground reaction force across multiple segments.

Some of the kinematic differences that we observed between models were due to the specific segment definitions. The Ghent and Vogel models advance distally from the MT joint, to the midfoot, to the TMT joint, and then to the forefoot. In contrast, the Oxford and Milwaukee models place the MT joint immediately proximal to the forefoot segment. This means that there is a greater degree of freedom distally for models like Ghent and Vogel, which impacts the angle measured at the MT joint. The variation in the degrees of freedom of a model impacts the joint angles observed between segments and this can affect kinetic calculations like power, which rely on angular acceleration.

It is important to understand that the kinematic and kinetic measurements are fueled by the motion of each individual foot segment. As more segments are added to MSFMs, the biomechanical model relies on more assumptions about the motion of these small foot segments. So, beyond a certain point, there is not necessarily more value in having more segments. It is also unclear if surface marker-based motion capture methods can obtain sufficient detail to represent all these segments, and especially the relative motion between adjacent segments. As an example, the Ghent model implements a medial/lateral split within the forefoot. While we know that the foot probably does differ kinematically and kinetically along this split, it is difficult to be certain that the model is effectively capturing what would be very fine, small movements between the second and first metatarsals. Overall, there are limitations as MSFMs continue to increase in number of segments and detail, beginning with the ability of motion capture to track such small segments.

A primary limitation of this study is that we did not have a plantar pressure map to distribute the ground reaction force across multiple segments throughout the entire foot. Instead, we manually assigned the force vector to the forefoot segment in Visual 3D. Our models still accounted for the center of pressure location but probably overestimated forefoot moments during time periods where the heel contacted the ground. Future work will incorporate plantar pressure to assign the force vectors throughout the foot accordingly. Additionally, some of these models have only ever been used for kinematic measurements, and there were technical challenges associated with updating them for kinetic measurements. For example, Visual 3D transmits moments and forces between segments linearly and did not allow for transmission medially/laterally between the forefoot segments the Ghent Model. Also, the Milwaukee model uses X-ray imaging in conjunction with motion capture to evaluate foot segments and movement, a practice that was not implemented here. As a result, segment orientations may differ between our model and the published Milwaukee model, although the overall kinematic patterns should be preserved. Lastly, these models do not account for each individual MT or MTP joint and we expect that these results might differ for each joint.

## Conclusion

This study demonstrates that the Oxford Foot Model, Milwaukee Foot Model, Vogel Foot Model, and Ghent Foot Model do not provide the same results regarding angle, moment, and power of the tibiotalar, midtarsal, and metatarsophalangeal joints. The changes in segment definition in these models impact the degrees of freedom in ways that alter the measured kinematic function of the foot, which in turn impacts the kinetic results. This work suggests that as MSFMs differ from one another simply in their construction, the results obtained from the model will change. Thus, there is a need to standardize the design of MSFMs to allow for more valuable and widespread comparisons across studies. Future work should be conducted to better understand the accuracy of tracking the small segments of the feet and the impact this has on the biomechanical measurements provided by MSFMs.

## Acknowledgements

This research is funded by the NIH R15HD104169.

