## Appendices for "How many segments are enough to biomechanically model the feet? A comparison of inverse kinematics and dynamics in multisegmented foot models"

**Appendix A**

| **Oxford Segments** | **Oxford Marker Set** | **Our Marker Set** |
| --- | --- | --- |
| **Shank** | Tibial tuberosity  Head of fibula  Lateral shank  Anterior shin  Medial malleolus  Lateral malleolus | Medial knee  Lateral condyle tibia  Medial superior tibia  Lateral superior tibia  Medial inferior tibia  Lateral inferior tibia  Medial malleolus  Lateral malleolus |
| **Rearfoot** | Inferior heel  Superior heel  Posterior calcaneus (wand)  Sustentaculum tale  Lateral calcaneus | Posterior inferior calcaneus  Posterior superior calcaneus  Medial calcaneus  Lateral calcaneus |
| **Forefoot** | Base of 1^st^ metatarsal  Head of 1^st^ metatarsal  Base of 5^th^ metatarsal  Head of 5^th^ metatarsal | Medial distal head MTI  Medial proximal head MTI  Dorsal distal head MTI  Dorsal distal head MTII  Dorsal proximal head MTII  Lateral distal head MTV  Lateral proximal head MTV |
| **Hallux** | Base of hallux | Distal head of 1^st^ phalange  Medial side of 1^st^ phalange |

| **Milwaukee Segments** | **Milwaukee Marker Set** | **Our Marker Set** |
| --- | --- | --- |
| **Shank** | Medial superior anterior tibia  Medial malleolus  Lateral malleolus | Medial knee  Lateral condyle tibia  Medial superior tibia  Lateral superior tibia  Medial inferior tibia  Lateral inferior tibia  Medial malleolus  Lateral malleolus |
| **Rearfoot** | Calcaneal tuberosity  Medial calcaneus  Lateral calcaneus | Posterior inferior calcaneus  Posterior superior calcaneus  Medial calcaneus  Lateral calcaneus |
| **Forefoot** | Medial head of 1^st^ metatarsal  Tuberosity of 5^th^ metatarsal lateral  Lateral head of 5^th^ metatarsal | Cuboid  Navicular  Cuneiform  Medial proximal head MTI  Medial distal head MTI  Dorsal distal head MTI  Dorsal proximal head MTII  Dorsal distal head MTII  Lateral proximal head MTV  Lateral distal head MTV |
| **Hallux** | Hallux triad x-axis  Hallux triad y-axis  Hallux triad z-axis | Distal head of 1^st^ phalange  Medial side of phalange |

| **Ghent Segments** | **Ghent Marker Set** | **Our Marker Set** |
| --- | --- | --- |
| **Shank** | Medial condyle tibia  Fibula head  Tibial tuberosity  Midway medial side tibia  Lateral malleolus  Medial malleolus | Medial knee  Lateral condyle tibia  Medial superior tibia  Lateral superior tibia  Medial inferior tibia  Lateral inferior tibia  Medial malleolus  Lateral malleolus |
| **Rearfoot** | Posterior inferior calcaneus  Posterior superior calcaneus  Sustentaculum Tale  Tuberculum peronei | Posterior inferior calcaneus  Posterior superior calcaneus  Medial calcaneus  Lateral calcaneus |
| **Midfoot** | Cuboid  Navicular  Dorsum of foot | Cuboid  Navicular  Cuneiform |
| **Medial Forefoot** | Base metatarsal 1  Shaft metatarsal 1  Lateral side of 1^st^ metatarsal head | Medial proximal head MTI  Medial distal head MTI  Dorsal distal head MTI |
| **Lateral Forefoot** | Dorsal metatarsals 2-5 (triad plate)  Head metatarsal 2  Lateral side base of 5^th^ metatarsal  Lateral side head metatarsal 5 | Dorsal proximal head MTII  Dorsal distal head MTII  Lateral proximal head MTV  Lateral distal head MTV |
| **Hallux** | Distal end of hallux  Proximal end hallux  Shaft of hallux (rod) | Distal head of 1^st^ phalange  Medial side of phalange |

**Appendix B**

**Markers:**

| Segment | Marker Name | Anatomic Location |
| --- | --- | --- |
| Shank | MED_KNEE  LAT_CON_TIB  MED_SUP_TIB  LAT_SUP_TIB  MED_INF_TIB  LAT_INF_TIB  MED_MAL  LAT_MAL_FIB | Medial knee  Lateral condyle tibia  Medial superior tibia  Lateral superior tibia  Medial inferior tibia  Lateral inferior tibia  Medial malleolus  Lateral malleolus |
| Hindfoot | POST_INF_CALC  POST_SUP_CALC  MED_CALC  LAT_CALC | Posterior inferior calcaneus  Posterior superior calcaneus  Medial calcaneus  Lateral calcaneus |
| Midfoot | CUBOID  MED_NAV  MED_CUN | Cuboid  Medial navicular  Medial cuneiform |
| Forefoot | MTI_PROX  MTI_DIST  SUP_PHAL  MTII_PROX  MTII_DIST MTV_PROX  MTV_DIST | Medial proximal head MTI  Medial distal head MTI  Dorsal distal head MTI  Dorsal proximal head MTII  Dorsal distal head MTII  Lateral proximal head MTV  Lateral distal head MTV |
| Phalanges | BIG_TOE  MED_PHAL | Distal head of 1^st^ phalange  Medial side of phalange |

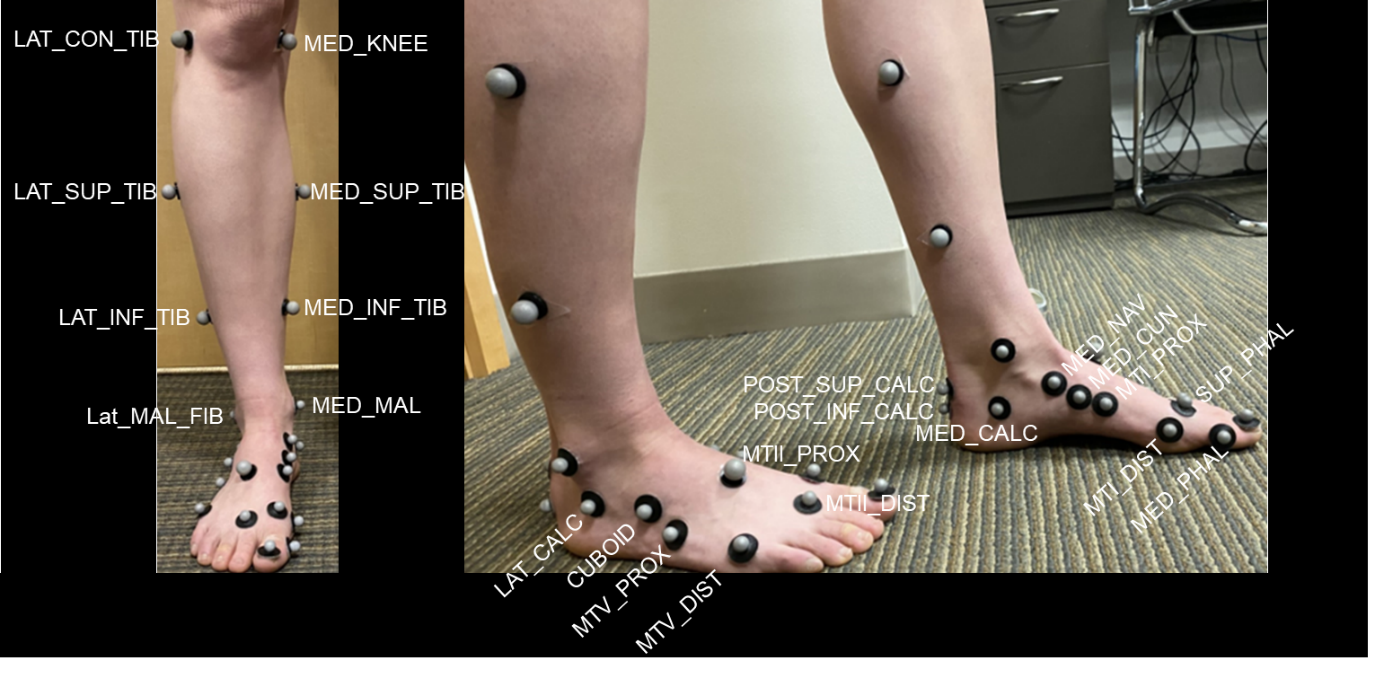

**Shank Segment**

Tracking Markers: R_LAT_INF_TIB, R_MED_INF_TIB, R_MED_MAL

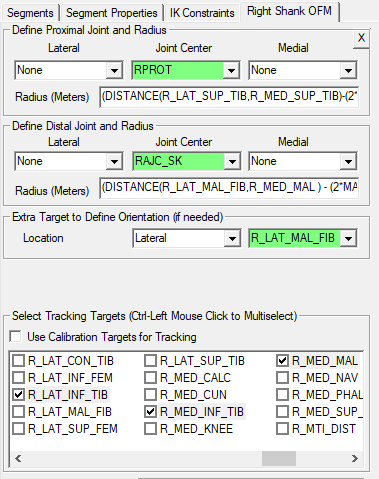

Radius (m): (DISTANCE(R_LAT_SUP_TIB,R_MED_SUP_TIB)-(2*MARKER_RADIUS))/2

Radius (m): (DISTANCE(R_LAT_SUP_MAL_FIB,R_MED_MAL)-(2*MARKER_RADIUS))/2

**Shank Segment Properties**

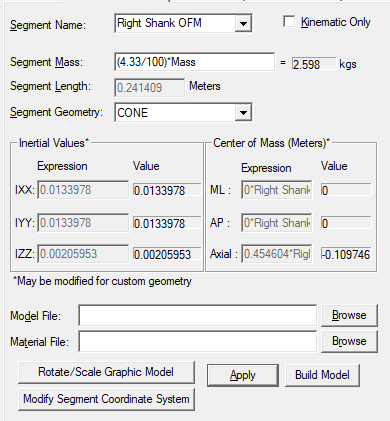

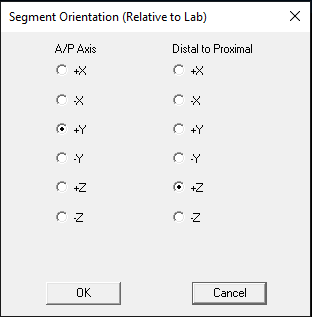

Segment masses were assigned based on known anthropometric values. Mass of the foot was divided among the segments based on the size of the segments.

**Shank Segment – Landmarks**

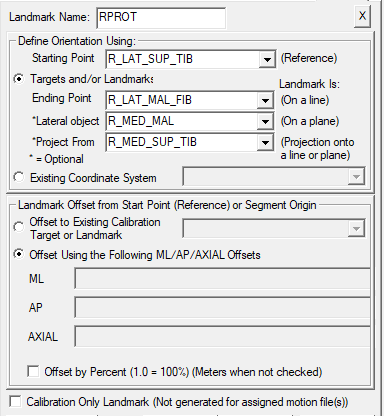

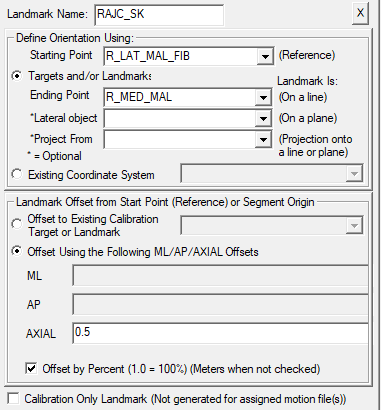

**Hindfoot Segment**

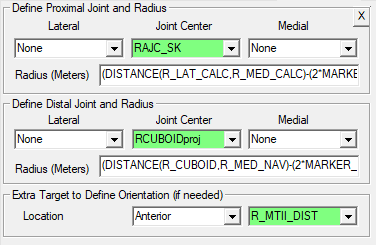

Radius (m): (DISTANCE(R_CUBOID,R_MED_NAV)-(2*MARKER_RADIUS))/2

Radius (m): (DISTANCE(R_LAT_CALC,R_MED_CALC)-(2*MARKER_RADIUS))/2

Tracking Markers:

R_LAT_CALC

R_MED_CALC

R_MED_CUN

**Rearfoot Segment Properties**

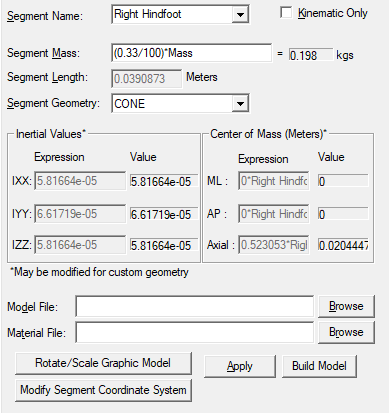

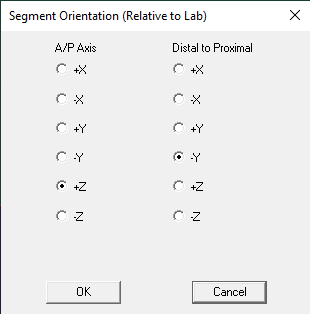

**Rearfoot Segment – Landmarks**

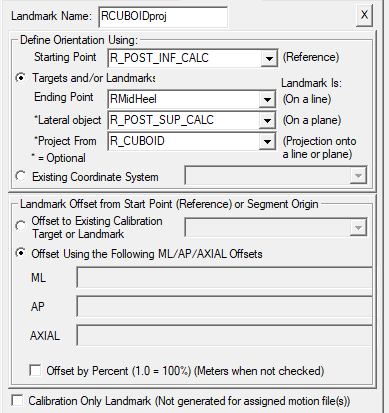

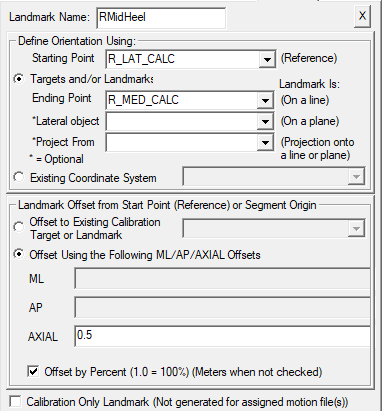

**Midfoot Segment**

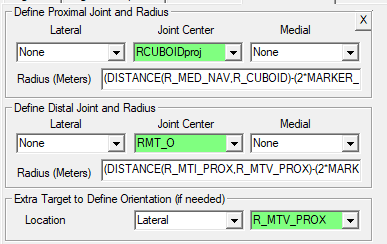

Radius (m): (DISTANCE(R_MED_NAV,R_CUBOID)-(2*MARKER_RADIUS))/2

Radius (m): (DISTANCE(R_MTI_PROX,R_MTV_PROX)-(2*MARKER_RADIUS))/2

Tracking Markers:

R_CUBOID

R_MED_CUN
R_MED_NAV

**Midfoot Segment Properties**

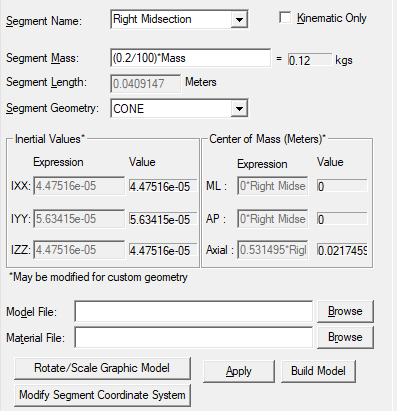

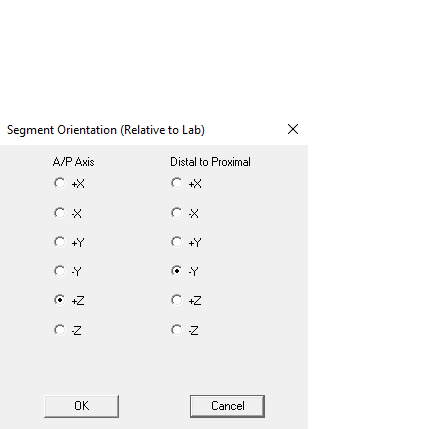

**Midfoot Segment – Landmarks**

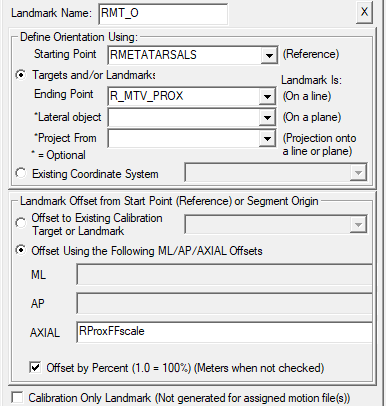

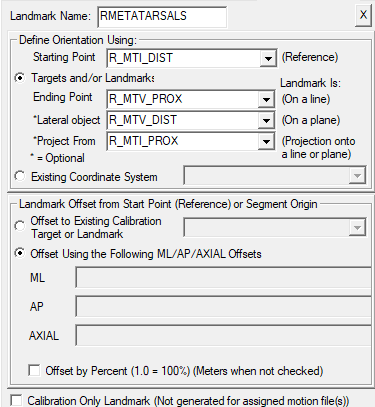

**Forefoot Segment**

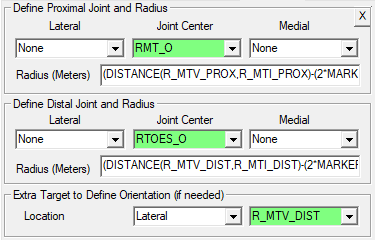

Radius (m): (DISTANCE(R_MTV_PROX,R_MTI_PROX)-(2*MARKER_RADIUS))/2

Radius (m): (DISTANCE(R_MTV_DIST,R_MTI_DIST)-(2*MARKER_RADIUS))/2

Tracking Markers:

R_MTI_DIST

R_MTII_PROX

R_MTV_DIST

R_MTV_PROX

**Forefoot Segment Properties**

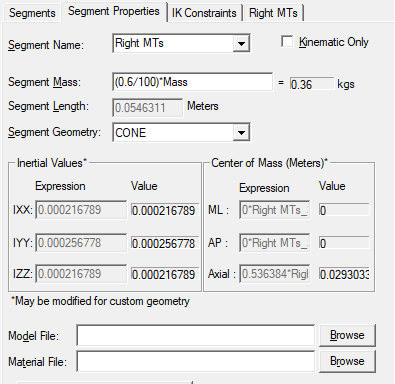

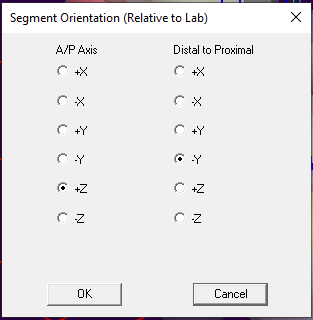

**Forefoot – Landmarks**

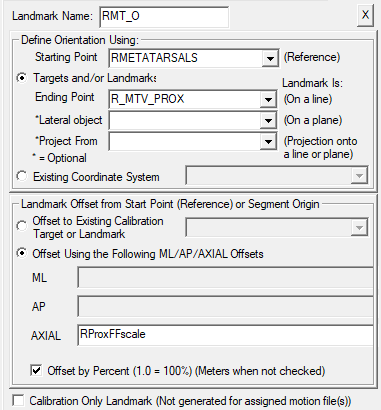

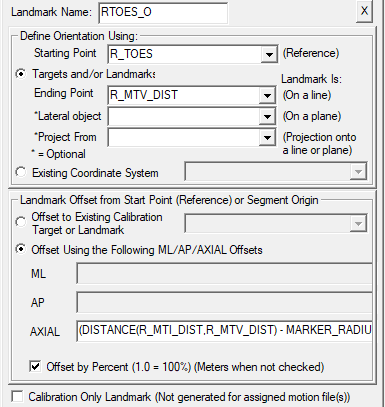

AXIAL: (DISTANCE(R_MTI_DIST,R_MTV_DIST) - MARKER_RADIUS) / (2*DISTANCE(R_MTI_DIST,R_MTV_DIST))

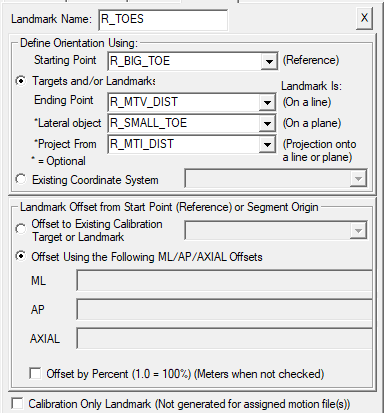

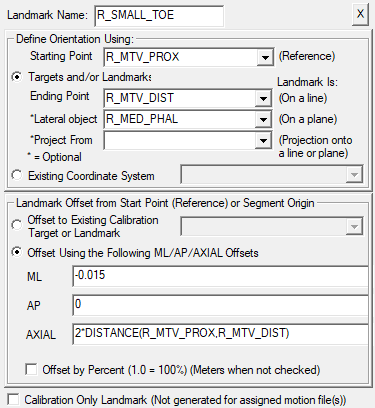

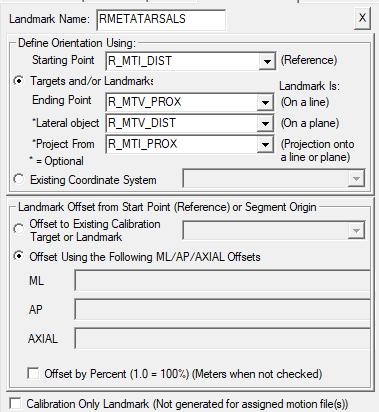

**Phalanges Segment**

Tracking Markers:

R_SMALL_TOE

R_MIDDLE_TOE

R_MTI_DIST

R_MTV_DIST

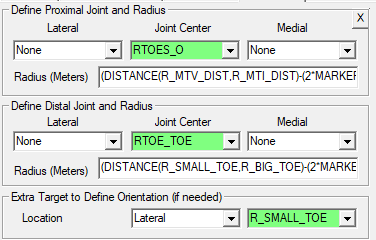

Radius (m): (DISTANCE(R_MTV_DIST,R_MTI_DIST)-(2*MARKER_RADIUS))/2

Radius (m): (DISTANCE(R_SMALL_TOE,R_BIG_TOE)-(2*MARKER_RADIUS))/2

**Phalanges Segment Properties**

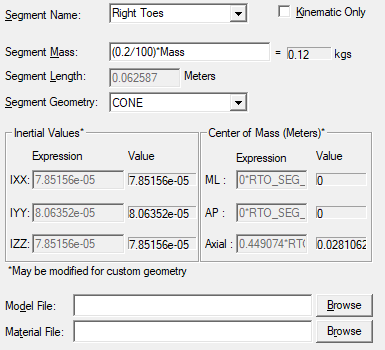

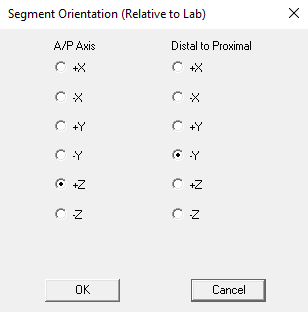

In order to incorporate a phalanges segment we used Visual 3D to construct landmarks for the middle toe and little toe based on the markers we had available. We recommend adding a reflective marker on the lateral side of the distal 5^th^ phalange and to the dorsal side of the distal 3^rd^ phalange during motion capture in place of the “SMALL_TOE” and “MIDDLETOE” landmarks.

**Phalanges Segment – Landmarks**

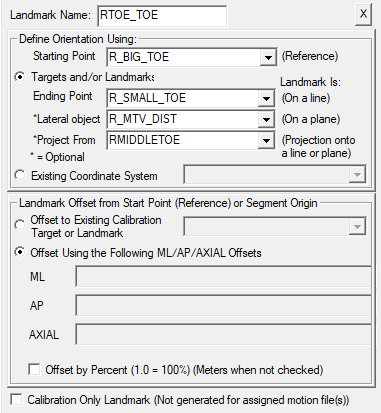

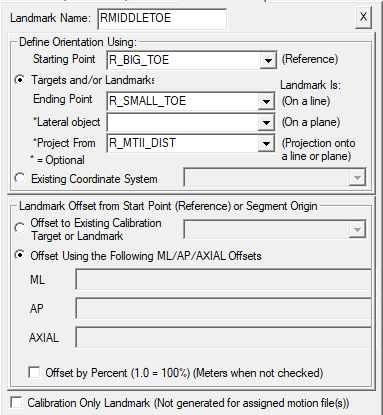
